# mtgRNA-db: An annotated database of multi-target CRISPR-Cas9 guide-RNAs in the human genome

**DOI:** 10.1101/2025.05.07.652656

**Authors:** Aram S. Modrek, Khoi Huynh, Catherine Do, Jerome Karp, Yingwen Ding, Ze-Yan Zhang, Ravesanker Ezhilarasan, Belen Valor, Giulia Cova, Jane A. Skok, Erik P. Sulman

**Author notes:** Contributed equally to this work (Jane A. Skok, Erik P. Sulman).

## Abstract

CRISPR-Cas9 guide-RNA design tools can be used to identify guide-RNAs that target single human genomic loci. However, these approaches limit their effect to a single locus. Here, we generate a database of potential individual guide-RNAs that target multiple sites in the genome at once. All guide-RNAs in this database are curated with on- and off-target quantification and enrichment scores for genomic elements such as gene elements, regulatory elements, repetitive elements, and transcription factor motifs. This tool enables rationale mass targeting of genomic loci with a single guide for functional studies. We created a web-app for user-friendly guide-RNA selection (https://modreklab.shinyapps.io/guiderna/).

## Introduction

Few strategies are available to target multiple genomic loci simultaneously using engineered tools in the same cell. Clustered Regularly Interspaced Short Palindromic Repeats and CRISPR-associated protein 9 (CRISPR-Cas9), Transcription Activator-Like Effector Nucleases (TALENs), and Zinc Finger Nucleases (ZFNs) are prime examples of programmable and sequence-specific molecules that can be engineered to target genomic loci(1), but current tools, databases or software do not allow them to be selected efficiently to target multiple loci at once, lack target annotation information of targets and may not be user friendly(2).

Multi-target guide-RNAs have been used experimentally to interrogate multiple loci at once. Ha and colleagues seeded CRISPR-Cas9 guide sequences from highly degenerate SINE (Short interspersed nuclear element) motifs, these guide RNAs were used to induce massive DNA-damage in parallel(3). Similar approaches were used by Fellman and colleagues who leveraged the presence of repeating and widespread mutational alterations in cancers with multi-target guide-RNAs to create DNA damage along many areas in the genome at once(4). These approaches employ guide-RNAs with highly degenerate sequences and do not provide users the flexibility to pick which genomic loci to methodically target. Other strategies for multi-loci genome targeting include non-programmable sequence-specific endonucleases that can be expressed in cells(5,6). The draw-back to this approach is that modifying the specificity of endonucleases is difficult, and nuclease activity can be blocked by DNA methylation(6,7).

Given the limitations in current strategies, we systematically searched for and compiled CRISPR-Cas9 guide-RNAs in the human genome and generated a database of curated guide-RNAs that can target 10 or more sites at once. CRISPR-Cas9 was used as the system of choice for several reasons: CRISPR-Cas9 is a robust well-characterized tool, it is widely used by biologists for a variety of applications, it does not require specialized tools or reagents to use, and the Cas9 trans-gene can be modified to create nicks, or be inactive and be conjugated to other effector payloads(8), providing a high level of flexibility. Therefore, we believe this database will serve as a resource to provide researchers with a curated list of multi-target human CRISPR-Cas9 guide-RNAs.

## Materials and Methods

All analysis and visualization was performed using R version 4.1.2 on a high-performance linux-based computer cluster. CRISPR-Cas9 PAM definitions and guide RNA sequences from PAM sites were obtained from (9) using the hg38 (GRCh38) reference genome. Cas-OFFinder (10,11) was used to calculate on-target and off-target matches for every possible guide-RNA. Aggregated results were combined, and quality controlled, as outlined in the results section, using the tidyverse R suite of tools. Genomic enrichment analysis was performed using genetic annotation libraries: ENCODE cis-regulatory elements(12), GENCODE (version 41) gene elements (13), 56 repetitive elements(14), and 340 transcription factor binding motifs from ENCODE ChIP-seq data(12,15-19). To perform a fisher’s exact test for genomic enrichment analysis, we used Locus OverLap Analysis (LOLA) (20). Web-application for user-friendly resource mining was developed using R-shiny app.

## Results

The pipeline to create mtgRNA-db is outlined in **Figure 1A**. The goal of the workflow was to identify all human CRISPR-spCas9-mtgRNAs that have 10 or more on-targets to the human genome. The starting point for database creation was all possible CRISPR-spCas9 sites in the human genome, as defined by the presence of a 5’-NGG sequence protospacer adjustment motif (PAM) site on either strand, of which there were approximately 305 million(9) in hg38. We set each of these 305 million PAM sites as seed sequences and extended each one to generate 305 million 18-nucletodite guide-RNA sequences as a starting point (**Figure 1B**). Of note, we picked 18 nucleotides because it would theoretically maximize the number of guides that target multiple sites in the genome (as opposed to a longer 20 nucleotide guide RNA). We did not search for shorter sequences (fewer then 17 nucleotides) because prior studies suggest that shorter guide-RNAs may not allow wild-type-CRISPR-Cas9 to function properly(21).

**Figure 1:**
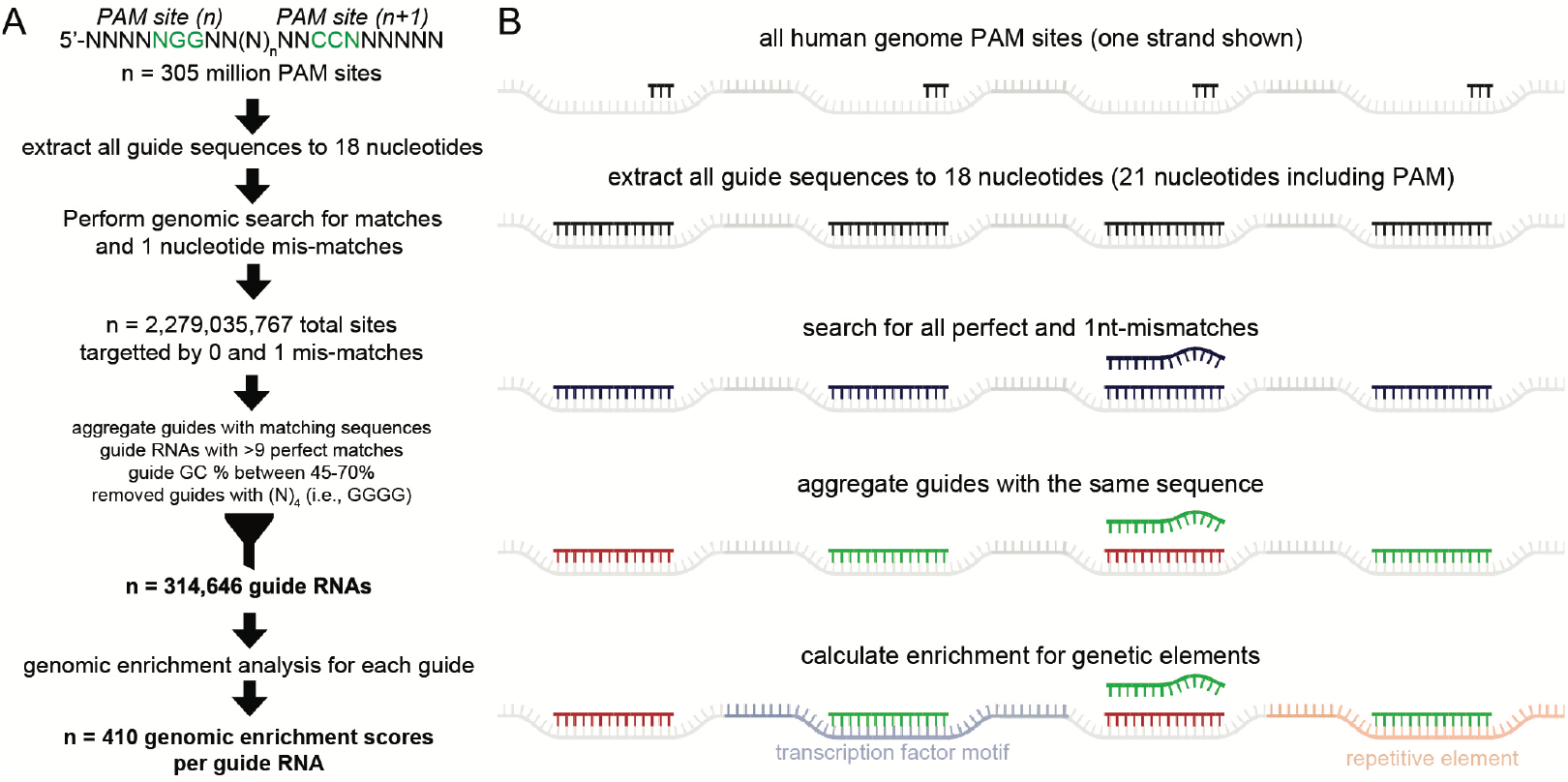
Generation of a human multi-target guide-RNA (mtgRNA) CRISPR-Cas9 database. **A)** Process for mtgRNA database generation. All human PAM sites were used as a seed sequence to generate 18-nucleotide guides. This library of guides was then consolidated for unique and matching loci, perfect match (0) and 1-nucleotide mismatches (off-targets) were quantified, and quality control measures were put into place. The ensuing guides were then subject to genomic enrichment analysis for further curation. **B)** Illustration of the pipeline in (A) used to generate the mtgRNA database. After all guide-RNAs were identified, guides were aggregated and consolidated to ascertain which guide-RNAs were in fact multi-target sequences. These same guides were then interrogated for enrichment of known genomic elements.

With the ∼305 million sequences generated, we took advantage of the finding that some of these guide-RNA sequences are the same, meaning that they also could target more than one site in the genome. After aggregating guides with the same sequence, each individual guide was subject to on-target quantification, i.e., how many sites in the genome does it perfectly match to? This was performed using Cas-OFFinder(10,11). Potential off-target quantification was also performed; this is checked by looking for 1 nucleotide mismatch outside of the PAM sequence to understand how many additional sites in the genome it can match to. This calculation was only performed for zero (perfect match) and single nucleotide mismatches, generating ∼2.2×10^9^ total genomic hits by all the guide-RNAs. We did not go beyond 1-nucleotide mismatch for this database, as the number of alignments and computational time required grows exponentially. However, knowing the number of potential off-targets with a single nucleotide mismatch for a given guide can serve as a surrogate for picking candidate guide-RNAs for further individual interrogation (i.e., prediction of 2-or 3-nucleotide mismatches, RNA bulge, DNA bulge, and other statistics which require more computational effort and can be performed on individual guide-RNAs with existing tools such as Cas-OFFinder, Cas-Designer, and Cas-Analyzer)(10).

The final step of mtgRNA generation was sequence quality control. We kept guide-RNAs that targeted more than nine genomic loci and had GC percentages between 45% and 75% to avoid highly repetitive degenerate sequences. We also believe this would allow for easier subcloning into vectors since oligonucleotides with very low or high GC content are difficult to subclone. We removed guide-RNAs that had four or more single nucleotide repeats for similar reasons (i.e., GGGG).

These quality control steps yielded a total of 314,646 guide-RNAs which met our criteria for inclusion into the database and further annotation. Visualization of perfect genome matches versus the number of 1-nulceotide mismatches reveals two key characteristics of the mtgRNAs identified (**Figure 1A**). First, the number of guide-RNAs that target >1000 perfect genome sites are relatively rare. Second, the number of multi-target guides that have a low number of mismatches is also an important consideration as some guides may have ∼10-100 perfect genome matches, but have >10,000-100,000 single nucleotide mismatches. While the number of targets is a consideration to take in guide selection, where the guides target (genomic location and context) is equally as important.

In **Figure 2B** we show four instances of single mtgRNAs that have different chromosomal distributions, two which target single chromosomes, and two which target multiple chromosomes, for example. Although we annotate each guide-RNA with the number of genomic matches (perfect matches, 1-nucleotide mismatch, and number of chromosomes covered), knowing the genetic elements that are potentially targeted by each multi-target guide-RNA may also be a valuable tool to those looking to target multiple functional elements in the genome with a single guide-RNA. Conversely, it may be of experimental importance avoid systematically targeting a functional genomic element. Therefore, we collected widely used genetic annotation libraries (ENCODE cis-regulatory elements(12), GENCODE (version 41) gene elements(13), 56 repetitive elements(14), and 340 transcription factor binding motifs from ENCODE ChIP-seq data(12,15-19)). In total, there were 410 unique genomic elements to perform enrichment testing for, and each guide was tested individually against all elements.

**Figure 2:**
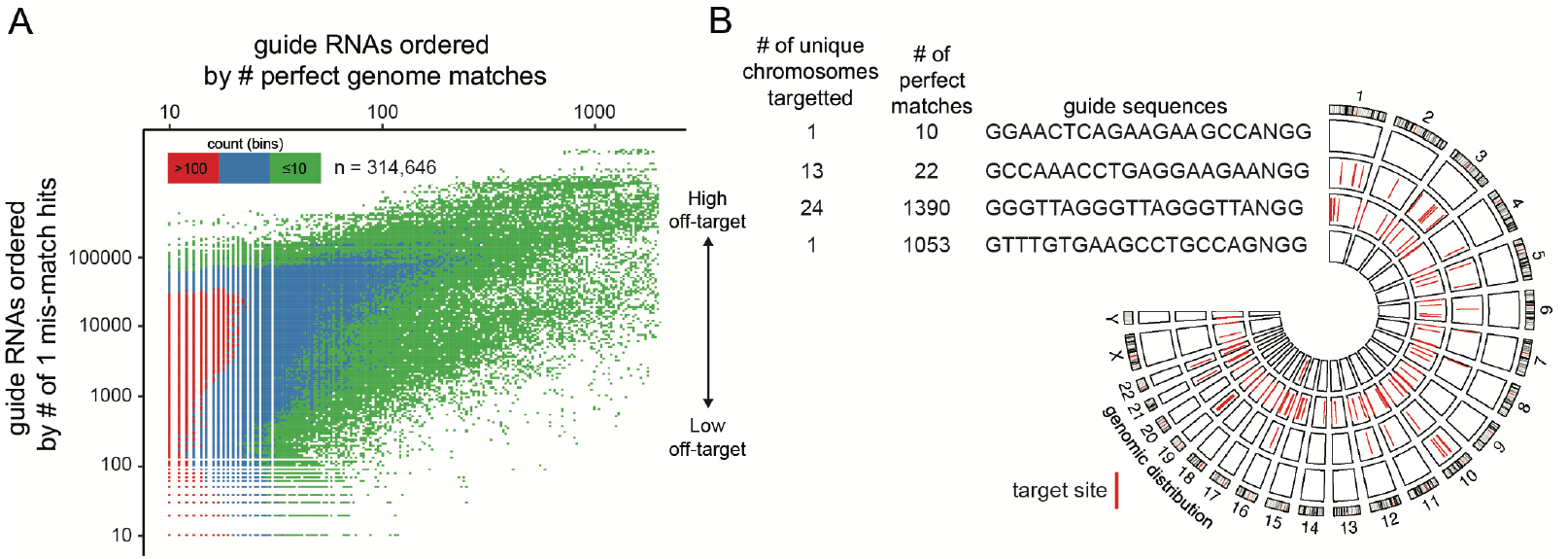
Multi-target guide-RNAs vary in their number of targeted genomic loci and chromosome positioning. **A)** A binned scatter plot depicting the number of perfect matches on the x-axis versus the number of 1-nucleotide mismatches on the y-axis. The number of binned counts represents the number of guide-RNAs that fall into the match counts as defined by the axes. This shows that most mtgRNAs have <100 perfect matches to the genome and high potential for off-targeting (>1000 1-nt mismatches). **B)** Four example guide-RNAs which show unique chromosome distributions and genome matches, illustrating the diversity in targeting potential.

To perform enrichment testing, we performed genomic enrichment analysis using a Locus OverLap Analysis (LOLA), a tool developed by Sheffield and Bock(20). This method uses a Fisher’s exact test with false discovery rate correction to see if any element in the genome is statistically enriched over other parts of the genome in a pairwise manner. We caution users that enrichment is not synonymous with overlap, meaning high enrichment of one particular genomic element does not exclude that the guide-RNA may be overlapping with other genomic elements. By using this method, a *P*-value, log odds ratio and number of overlaps can be assessed for each genomic element.

The aggregated result of this genomic enrichment analysis is visualized in **Figure 3**. The results are grouped by genomic element class: cis-regulatory elements (**Figure 3A**), gene-elements (**Figure 3B**), repetitive elements (**Figure 3C**), transcription factor binding motifs (**Figure 3D**). The results suggest that some genomic elements can be highly enriched for by single guide-RNAs (enrichment *P*-values on the x-axis), but many of these guide-RNAs have varying overlap proportions with individual genomic elements (% of guide hitting the genomic element), which could be an important experimental consideration when selecting a guide-RNA to use. We show individual examples of these guide-RNAs in **Figure 4**, including a guide-RNA enriched for exons (**Figure 4A**), and one enriched for repetitive elements (**Figure 4B**). We also show a guide-RNA that is enriched for a transcription factor binding motif (**Figure 4C**); as expected, we noticed a high degree of DNA motif similarity between the transcription factor binding motif and the multi-target guide-RNA.

**Figure 3:**
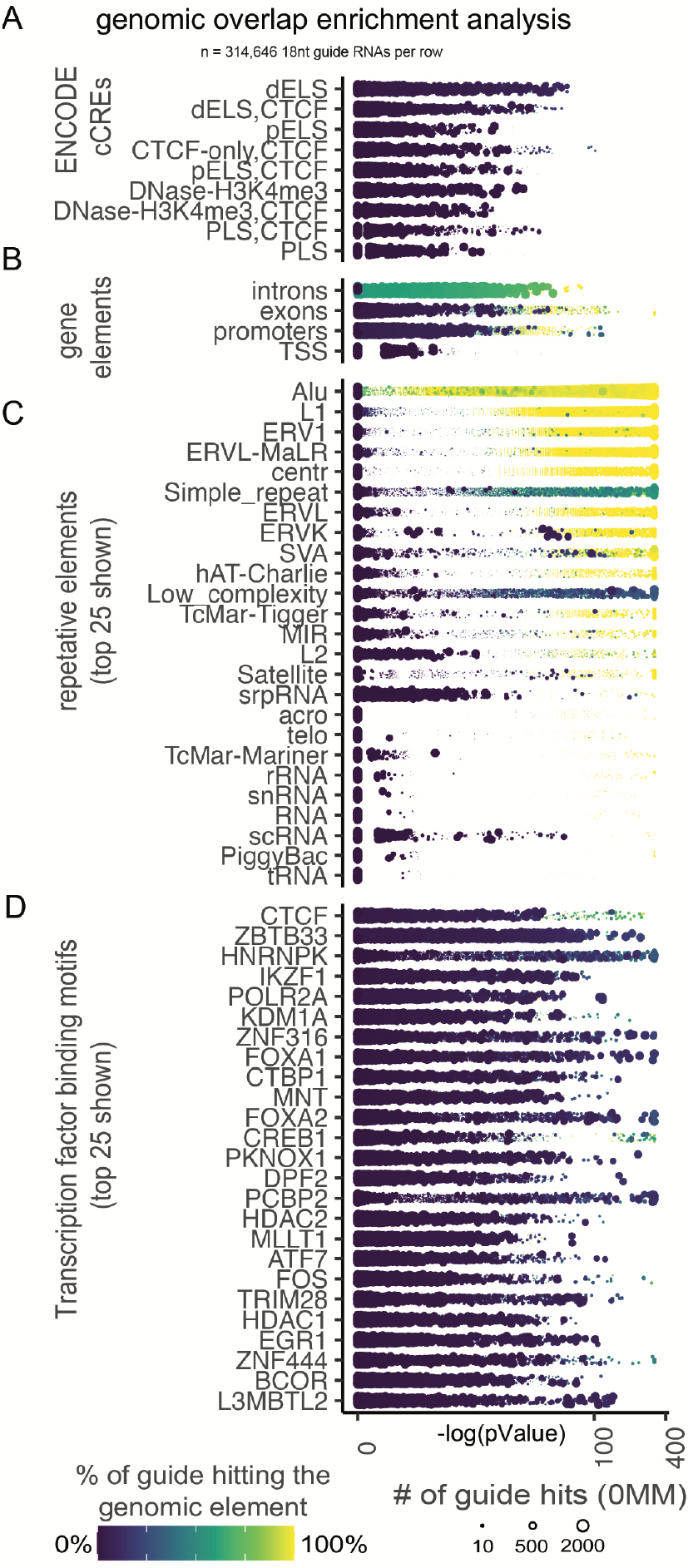
Genomic overlap enrichment analysis of mtgRNAs. Genomic enrichment analysis was performed for each guide-RNA. Each row has 314,646 data points. The datapoints are colored based on the percentage of the guide that hits the genomic element on that row. The size of each datapoint represents the total number of on-target hits and the position on the x-axis represents the *P*-value measure of how enriched it is for that element. Due to space limitations, the only the top 25 repetitive elements and top 25 transcription factor binding sites are shown. **A)**ENCODE cis-regulatory elements (cCREs). **B)**gene elements as defined by GENCODE. **C)** Repetitive elements **D)** Transcription factor binding motifs.

**Figure 4:**
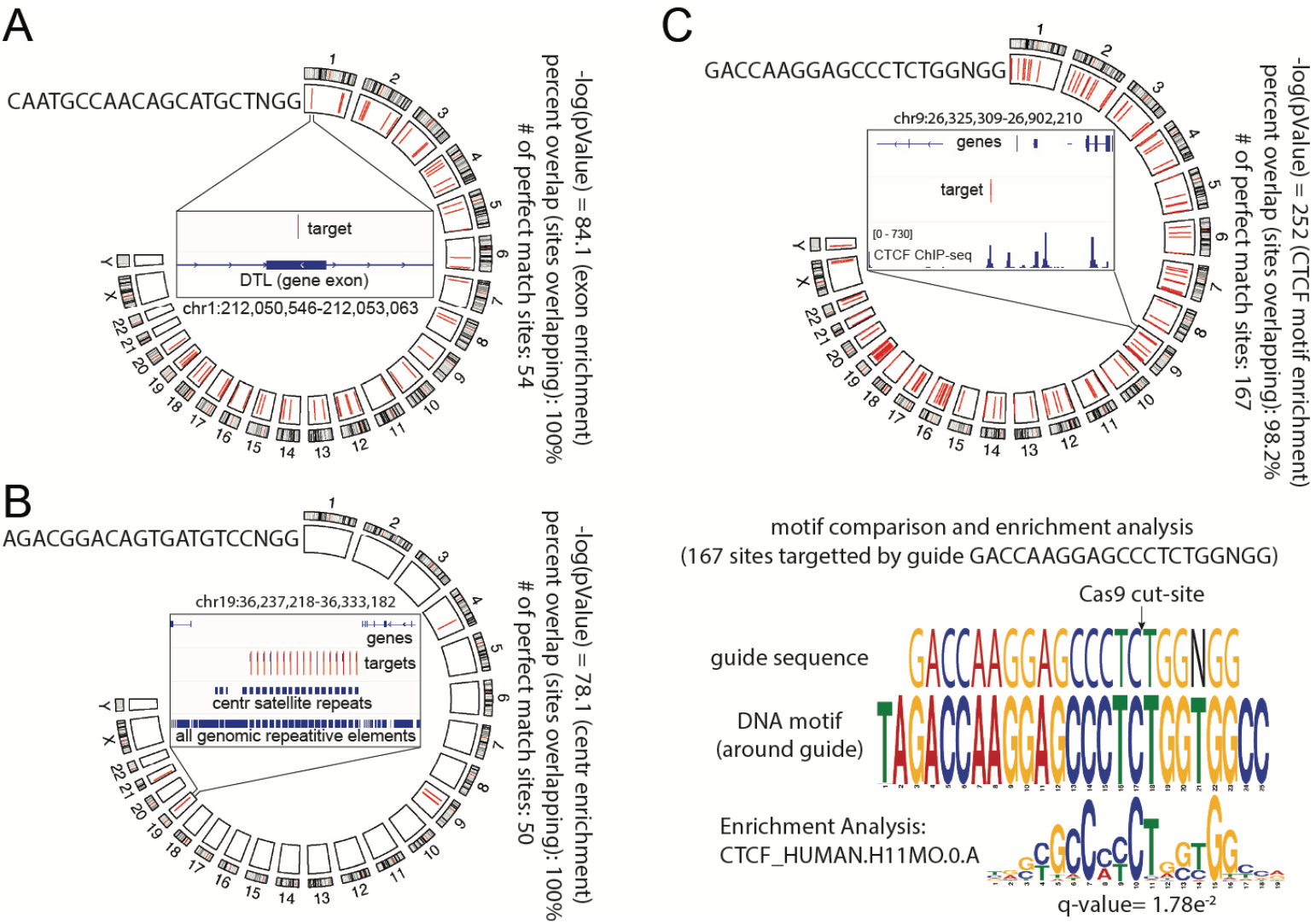
Target distributions of three mtgRNAs enriched for genomic elements. **A)** gene element enriched mtgRNA with enrichment for exons (100% of guide target sites, n=54, have exon overlap). Inset shows overlap on genome view. **B)** Repetitive element enriched mtgRNA with enrichment for centr satellite repeats (100% of guide target sites, n=50, have centr satellite repeat overlap). Inset shows overlap on genome view. **C)** transcription factor motif enriched mtgRNA with enrichment for CTCF binding motifs (98.2% of guide target sites, n=167, have CTCF motif overlap). Inset shows overlap with certain CTCF ChIP-seq peaks. Bottom panel shows a motif comparison of the guide sequence motif, the DNA around the guide, and the CTCF motif (q-value represents the statistical measure of the CTCF human DNA motif(22,23)).

We created a web-app for user-friendly mtgRNA selection and download (https://modreklab.shinyapps.io/guiderna/). Once suitable mtgRNAs are identified, we recommend empirically testing the number of guide-RNA targets first using bioinformatic verification followed by *in vitro* or *in vivo* verification.

## Discussion

There are open challenges with inducing DNA damage to study DNA damage repair and response(24). Current endonuclease systems used to study genome-wide DNA damage are limited by a fixed genomic target or targets, making them inflexible. In addition, many endonucleases have their activity blocked by DNA methylation of their cut site(6,7). On the other hand, current CRISPR-Cas9 guide-RNA design tools are designed to select and filter for guides that have a single on-target guide sequence and minimal off-targets. This database represents all the spCas9 guides which target many sites in the genome simultaneously. Therefore, investigators using this database can select single guides that target many areas of the genome, or single chromosomes, etc. for their experimental purposes. Multiple mtgRNAs may also be selected to use in screens (i.e., one guide-RNA per cell in a screen) or expressed in the same cell for combined use.

As an example, this database can be used to find a guide that targets multiple chromosomes at ten different locations, or a user could select a mtgRNA that targets thousands of sites along a single chromosome (**Figure 2B**). The utility of this depends on the unique biological questions and aims of the user. For those interested in creating pre-mapped DNA damage, this system could be used to study double-strand break biology, single-strand break biology (with a modified Cas9 enzyme) or bring Cas9 conjugated molecules to multiple sites at once. Due to the fact that damage is pre-mapped, the ability to study translocations and the genesis of other large structural variations may be enhanced. As another example, this technology could be used to achieve chromosome “painting” with deadCas9 conjugated to a fluorophore.

There are limitations to using these mtgRNAs for CRISPR-Cas9 targeting. Currently, this database is limited to 18 nucleotide guide-RNAs and uses the PAM sequence that is used by spCas9 (5’-NGG). Therefore, specific patterns of genome-wide targeting from a single guide may not be possible. Other limitations are the extent of off-target prediction which is limited to one nucleotide mismatch in this database (theoretically, 2 or more mismatches are possible, as well as RNA or DNA bulging). In the future, additional RNA-guided endonuclease systems may be integrated into this database to expand its use.

Within coding and non-coding regions of the genome, there are many different genomic elements which play distinct roles, many of which are actively being investigated. For example, cis-regulatory elements such as enhancers are responsible for regulating neighboring genes. Transcription factor (TF) binding motifs serve to act as binding sites for transcription factors. Although most of our genome is annotated by these distinct elements (repetitive elements, genes, TF binding sites and cis-regulatory elements), their collective function can be difficult to study as there is redundancy and sequence degeneracy along the genome. This can result in hundreds to thousands of binding sites for a single transcription factor, for example. Because of this high level of redundancy and degeneracy, our genome contains many elements that repeat themselves and contain similar sequences. This database takes advantage of this degeneracy by identifying and annotating mtgRNAs that are enriched for these genomic elements via exact matches.

To understand which mtgRNAs are enriched for known genomic elements, we took all 314,646 mtgRNAs and ran an enrichment analysis against aforementioned genomic elements, as stated in the construction and content section. This resulted in each mtgRNA being annotated with an enrichment score for each element (**Figure 3,4**). This method has its limitations. MtgRNAs with high enrichment scores for a particular element may still be enriched for other elements or have a high degree of targets along the genome that are not marked by known elements. We recommend screening candidate mtgRNAs for all enrichment scores and potential on-and off-targets prior to *in vitro* or *in vivo* testing.

In conclusion, we present mtgRNA-db, a curated resource of multi-target guide-RNAs for use with CRISPR-Cas9 in the human genome. Each individual mtgRNA is annotated with the number of target sites, off-target sites, number of chromosomes targeted, and enrichment scores for genomic elements. Future work and directions may include expanding this database to include other programmable nucleases and genomes from other organisms. We suggest that this database will serve as a useful tool for genome-wide functional studies using multi-target guide-RNAs to enable massively parallel targeting of the human genome.

## Authors’ contributions

ASM, JAS and EPS conceived the study. JAS and EPS supervised the project. ASM, KH, CD, JP and BV processed the raw data. AM, YD, ZYZ, RE and GC visualized the data and created figures. KH and JP created the web-application. ASM wrote the initial manuscript draft, and all authors provided critical reading and edits. All authors read and approved the final manuscript.

## Availability of data and materials

This resource can be downloaded and browsed on: https://modreklab.shinyapps.io/guiderna

## Declarations

Ethics approval and consent to participate: not applicable.

## Competing interests

The authors declare that they have no competing interests.

## Funding

This work was supported by the following funding sources: NIH 2R35GM122515, NIH 7K08CA263302, NYU NIH/NCATS UL1TR001445, American Brain Tumor

Association, V Foundation, Baxter Foundation, and Radiological Society of North America.

## Notes

https://modreklab.shinyapps.io/guiderna/

### Competing Interest Statement

The authors have declared no competing interest.

## References

1. Gaj, T., Gersbach, C.A. and Barbas, C.F., 3rd. (2013) ZFN, TALEN, and CRISPR/Cas-based methods for genome engineering. Trends Biotechnol, 31, 397–405.

2. Veluchamy, A., Teles, K. and Fischle, W. (2023) CRISPR-broad: combined design of multi-targeting gRNAs and broad, multiplex target finding. Sci Rep, 13, 19717.

3. Zou, R.S., Marin-Gonzalez, A., Liu, Y., Liu, H.B., Shen, L., Dveirin, R.K., Luo, J.X.J., Kalhor, R. and Ha, T. (2022) Massively parallel genomic perturbations with multi-target CRISPR interrogates Cas9 activity and DNA repair at endogenous sites. Nat Cell Biol, 24, 1433–1444.

4. Tan, I.L., Perez, A.R., Lew, R.J., Sun, X., Baldwin, A., Zhu, Y.K., Shah, M.M., Berger, M.S., Doudna, J.A. and Fellmann, C. (2023) Targeting the non-coding genome and temozolomide signature enables CRISPR-mediated glioma oncolysis. Cell Rep, 42, 113339.

5. Monnat, R.J., Jr., Hackmann, A.F. and Cantrell, M.A. (1999) Generation of highly site-specific DNA double-strand breaks in human cells by the homing endonucleases I-PpoI and I-CreI. Biochem Biophys Res Commun, 255, 88–93.

6. Iacovoni, J.S., Caron, P., Lassadi, I., Nicolas, E., Massip, L., Trouche, D. and Legube, G. (2010) High-resolution profiling of gammaH2AX around DNA double strand breaks in the mammalian genome. EMBO J, 29, 1446–1457.

7. Aymard, F., Bugler, B., Schmidt, C.K., Guillou, E., Caron, P., Briois, S., Iacovoni, J.S., Daburon, V., Miller, K.M., Jackson, S.P. and Legube, G. (2014) Transcriptionally active chromatin recruits homologous recombination at DNA double-strand breaks. Nat Struct Mol Biol, 21, 366–374.

8. Pulecio, J., Verma, N., Mejia-Ramirez, E., Huangfu, D. and Raya, A. (2017) CRISPR/Cas9-Based Engineering of the Epigenome. Cell Stem Cell, 21, 431–447.

9. Haeussler, M., Schonig, K., Eckert, H., Eschstruth, A., Mianne, J., Renaud, J.B., Schneider-Maunoury, S., Shkumatava, A., Teboul, L., Kent, J. et al. (2016) Evaluation of off-target and on-target scoring algorithms and integration into the guide RNA selection tool CRISPOR. Genome Biol, 17, 148.

10. Hwang, G.H., Kim, J.S. and Bae, S. (2021) Web-Based CRISPR Toolkits: Cas-OFFinder, Cas-Designer, and Cas-Analyzer. Methods Mol Biol, 2162, 23–33.

11. Bae, S., Park, J. and Kim, J.S. (2014) Cas-OFFinder: a fast and versatile algorithm that searches for potential off-target sites of Cas9 RNA-guided endonucleases. Bioinformatics, 30, 1473–1475.

12. Consortium, E.P. (2012) An integrated encyclopedia of DNA elements in the human genome. Nature, 489, 57–74.

13. Frankish, A., Diekhans, M., Jungreis, I., Lagarde, J., Loveland, J.E., Mudge, J.M., Sisu, C., Wright, J.C., Armstrong, J., Barnes, I. et al. (2021) Gencode 2021. Nucleic Acids Res, 49, D916–D923.

14. Jurka, J. (2000) Repbase update: a database and an electronic journal of repetitive elements. Trends Genet, 16, 418–420.

15. Consortium, E.P. (2011) A user’s guide to the encyclopedia of DNA elements (ENCODE). PLoS Biol, 9, e1001046.

16. Sloan, C.A., Chan, E.T., Davidson, J.M., Malladi, V.S., Strattan, J.S., Hitz, B.C., Gabdank, I., Narayanan, A.K., Ho, M., Lee, B.T. et al. (2016) ENCODE data at the ENCODE portal. Nucleic Acids Res, 44, D726–732.

17. Gerstein, M.B., Kundaje, A., Hariharan, M., Landt, S.G., Yan, K.K., Cheng, C., Mu, X.J., Khurana, E., Rozowsky, J., Alexander, R. et al. (2012) Architecture of the human regulatory network derived from ENCODE data. Nature, 489, 91–100.

18. Wang, J., Zhuang, J., Iyer, S., Lin, X., Whitfield, T.W., Greven, M.C., Pierce, B.G., Dong, X., Kundaje, A., Cheng, Y. et al. (2012) Sequence features and chromatin structure around the genomic regions bound by 119 human transcription factors. Genome Res, 22, 1798–1812.

19. Wang, J., Zhuang, J., Iyer, S., Lin, X.Y., Greven, M.C., Kim, B.H., Moore, J., Pierce, B.G., Dong, X., Virgil, D. et al. (2013) Factorbook.org: a Wiki-based database for transcription factor-binding data generated by the ENCODE consortium. Nucleic Acids Res, 41, D171–176.

20. Sheffield, N.C. and Bock, C. (2016) LOLA: enrichment analysis for genomic region sets and regulatory elements in R and Bioconductor. Bioinformatics, 32, 587–589.

21. Fu, Y., Sander, J.D., Reyon, D., Cascio, V.M. and Joung, J.K. (2014) Improving CRISPR-Cas nuclease specificity using truncated guide RNAs. Nat Biotechnol, 32, 279–284.

22. Bailey, T.L., Johnson, J., Grant, C.E. and Noble, W.S. (2015) The MEME Suite. Nucleic Acids Res, 43, W39–49.

23. Bailey, T.L., Boden, M., Buske, F.A., Frith, M., Grant, C.E., Clementi, L., Ren, J., Li, W.W. and Noble, W.S. (2009) MEME SUITE: tools for motif discovery and searching. Nucleic Acids Res, 37, W202–208.

24. Vitor, A.C., Huertas, P., Legube, G. and de Almeida, S.F. (2020) Studying DNA Double-Strand Break Repair: An Ever-Growing Toolbox. Front Mol Biosci, 7, 24.

